# Dose-dependent interferon programs in myeloid cells after mRNA and adenovirus COVID-19 vaccination

**DOI:** 10.1101/2025.08.15.668720

**Authors:** Giray Naim Eryilmaz, Yilmaz Yucehan Yazici, Radu Marches, Eleni P. Mimitou, Lisa Kenyon-Pesce, Kim Handrjek, Sonia Jangra, Michael Schotsaert, Adolfo García-Sastre, George A. Kuchel, Jacques Banchereau, Duygu Ucar

## Abstract

The SARS-CoV-2 pandemic provided a rare opportunity to study how human immune responses develop to a novel viral antigen delivered through different vaccine platforms. However, to date, no study has directly compared immune responses to all three FDA-approved COVID-19 vaccines at single-cell multi-omic resolution. We longitudinally profiled SARS-CoV-2–naïve adults (n=31) vaccinated with BNT162b2, mRNA-1273, or Ad26.COV2.S, integrating plasma cytokines, antibody titers, and single-cell multi-omic data (DOGMA-seq). We discovered a distinct, transient interferon program (ISG-dim) that emerged specifically 1-2 days after the first mRNA dose in ∼10% of myeloid cells. This state was characterized by ISGF3 complex activation and its target genes (e.g., *MX1*, *MX2*, *DDX58*), with transcriptional and epigenetic profiles distinct from the robust interferon program observed after mRNA boosting or a single Ad26.COV2.S dose (ISG-high). In vitro stimulation of human monocytes showed that IFN-α alone recapitulates ISG-dim, whereas both IFN-α and IFN-γ are required for ISG-high. These findings define dose-dependent interferon programming in human myeloid cells, highlight mechanistic differences between priming and boosting, with implications for optimizing vaccine platform choice, dose scheduling, and formulation.

## Introduction

The SARS-CoV-2 pandemic posed unprecedented global health challenges but also created a rare scientific opportunity to study how human immune responses develop to a novel viral antigen introduced through different vaccine platforms. Among these, the novel and highly effective mRNA vaccines BNT162b2 (Pfizer) and mRNA-1273 (Moderna) utilized lipid nanoparticles (LNPs) to deliver nucleoside-modified RNA encoding the SARS-CoV-2 spike (S) protein^1–4^. An alternative platform, Ad26.COV2.S (Johnson & Johnson), delivered spike-encoding DNA *via* a non-replicating adenoviral vector^5^. Although all three vaccines target the same antigen, their distinct formulations and platforms provide a unique framework to dissect human immune priming and boosting. Despite this opportunity, to our knowledge, no prior study has performed a head-to-head, single-cell multi-omic comparison of all three FDA-approved COVID-19 vaccines in SARS-CoV-2–naïve individuals.

Immune responses, particularly adaptive immune responses following the second dose of mRNA vaccines, have been well characterized, including the robust expansion of CD4 and CD8 T cells, memory B cells, and the production of SARS-CoV-2-neutralizing antibodies^6–15^. In contrast, early innate responses, especially in human myeloid cells following the first dose, remain poorly understood. While some studies noted interferon-stimulated gene (ISG) induction^13,16–18^, the magnitude, timing, and cell-type specificity of this response have not been resolved. Our study fills this critical gap by providing the first single-cell, multi-omic map of early innate priming in humans following mRNA vaccination.

Here, we leveraged the unique conditions of the early days of the pandemic to study how immune responses develop in SARS-CoV-2–naïve healthy adults (n = 31) vaccinated with either mRNA (BNT162b2^3^, mRNA-1273^4^) or the adenoviral vector (Ad26.COV2.S^5^) COVID-19 vaccines. Using single-cell multi-omics alongside cytokine and antibody profiling, we mapped the temporal dynamics of innate responses at single-cell resolution. We uncovered a distinct myeloid priming program induced by Type I interferons after the first mRNA dose, and a broader boosting response involving both Type I and Type II interferons after the second mRNA dose or Ad26.COV2.S. By defining dose- and vaccine-platform-specific interferon programs in human myeloid cells, our study provides a mechanistic framework for optimizing vaccine platform choice, scheduling, and formulation, with implications for improving protection in vulnerable populations.

## Results

### Head-to-head comparison of three COVID-19 vaccines in SARS-CoV-2–naïve adults

Thirty-one healthy adults with no prior history of SARS-CoV-2 infection or vaccination were enrolled between spring and summer 2021 at the UConn Health Center. Of these, twenty-eight participants received mRNA vaccines, including both BNT162b2 (n=22) and mRNA-1273 (n=6), while 3 received the adenoviral vector vaccine Ad26.COV2.S (Fig. 1A, Table S1). Age distributions were comparable across groups (Fig. 1A, Table S1).

**Figure 1.**
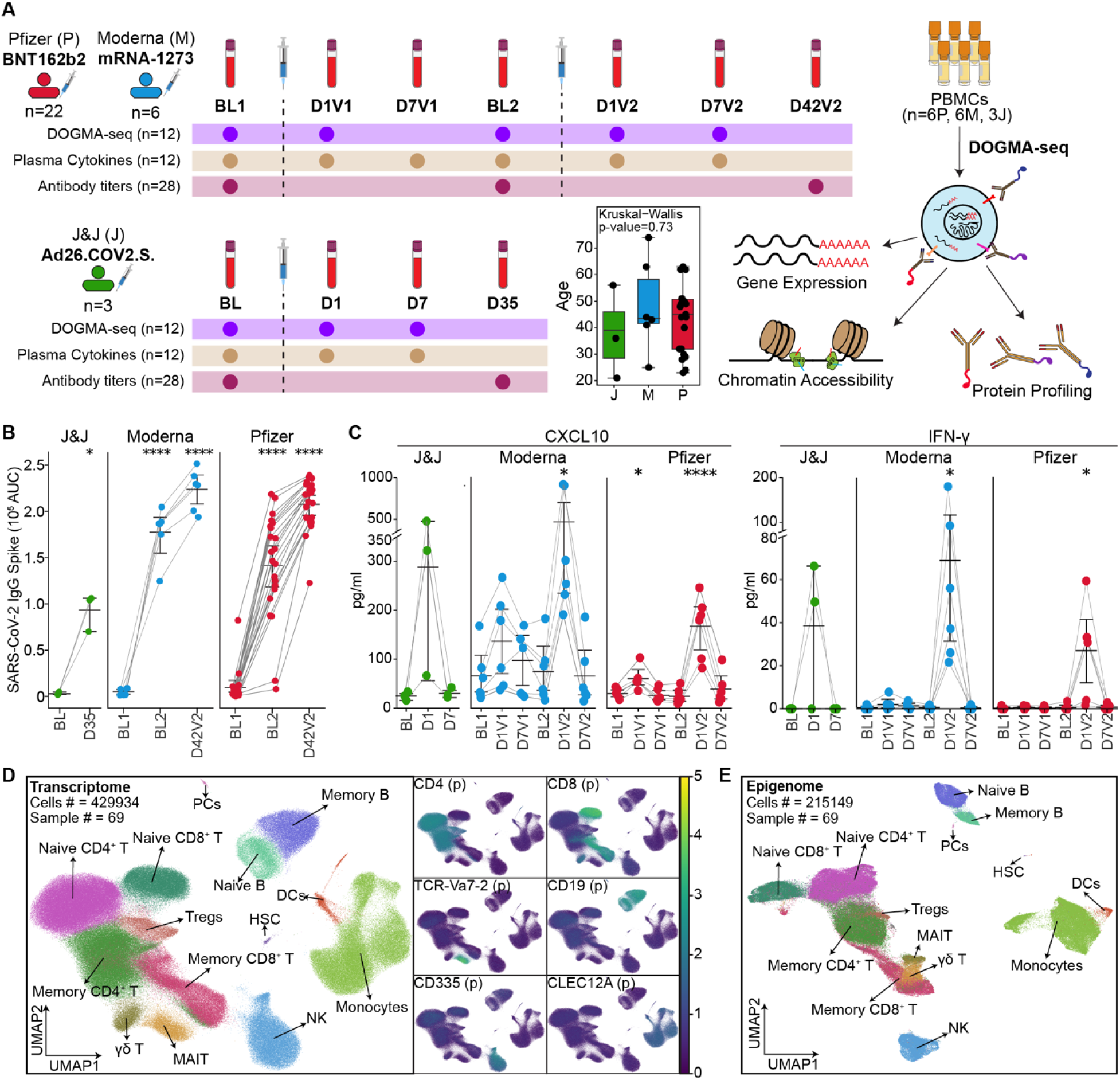
A longitudinal systems immunology study of immune responses to COVID-19 vaccines. **(A)** Study design. Blood samples were collected longitudinally from individuals (n = 31) vaccinated with either two doses of mRNA vaccines (Pfizer-BioNTech, Moderna) or a single dose of the adenovirus-based Johnson & Johnson (J&J) vaccine. Antibody titers were measured before and 3–4 weeks after each vaccination. A subset of donors (6 Pfizer, 6 Moderna, 3 J&J) was selected for in-depth profiling, including serum cytokine analysis and DOGMA-seq. Age distributions among vaccine groups were compared using the Kruskal–Wallis test. **(B)** IgG titers against SARS-CoV-2 spike protein measured by ELISA before and after vaccination. Antibody titers further increased after the mRNA booster dose. **(C)** Serum levels of CXCL10 and IFN-γ at key time points were quantified using ELLA. Both cytokines increased on day 1 (D1) after the adenoviral vaccine and following the mRNA booster dose. **(D–E)** UMAP projections of DOGMA-seq data for transcriptome (D, scRNA-seq) and chromatin accessibility (E, scATAC-seq) modalities. Canonical lineage markers (surface protein and gene expression) annotated major immune cell populations. **(B), (C)** Statistical significance was assessed using two-tailed paired t-tests: \**P* < 0.05, \*\**P* < 0.01, \*\*\**P* < 0.001, \*\*\*\**P* < 0.0001.

mRNA vaccines were administered in two doses, while Ad26.COV2.S was administered as a single dose by the CDC recommendations. Blood samples were collected at key timepoints: baseline (pre-vaccination), day 1 (innate responses), day 7 (early adaptive responses), and days 25–35 (antibody responses) after each vaccine dose (Fig. 1A, Table S1), totaling seven timepoints for mRNA vaccines and four timepoints for Ad26.COV2.S. The isolated peripheral blood mononuclear cells (PBMCs) were used for genomic profiling, and plasma samples were used for cytokine and antibody titer profiling.

Anti–SARS-CoV-2 spike protein IgG titers were measured by ELISA at baseline and 3–4 weeks after each vaccination. All vaccine groups showed a significant increase in antibody titers after the first dose (*p* < 0.0001), followed by an additional increase after the second mRNA dose (Fig. 1B). After two doses, individuals vaccinated with mRNA vaccines had significantly higher titers (p=0.0369) than those receiving a single dose of Ad26.COV2.S (Fig. S1A). Despite the dose difference between BNT162b2 (30 µg) and mRNA-1273 (100 µg), antibody titers were comparable between the two mRNA groups at both time points (Fig. 1B, Fig. S1A). No correlation was observed between antibody titer levels and participant age (Fig. S1B).

Plasma levels of six cytokines were quantified at baseline, day 1, and day 7 after each vaccination using the ELLA® platform (Fig 1C, Fig. S1C, and Table S2). The first dose of mRNA vaccines modestly elevated CXCL10 levels at day 1 (*p* = 0.14 for Moderna, *p* = 0.013 for Pfizer), while the second dose induced stronger induction of both CXCL10 and IFN-γ (CXCL10: *p* = 0.019 for Moderna, *p* = 0.00036 for Pfizer; IFN-γ: *p* = 0.019 for Moderna, *p* = 0.0125 for Pfizer, Fig. 1C). In contrast, both IFN-γ and CXCL10 levels increased to levels comparable to those induced by mRNA vaccine recipients at day 1 following Ad26.COV2.S. No significant changes were observed for IL-8, TNF-α, CCL2, or IL-10 levels following any vaccine (Fig. S1C).

Together, these data showed robust antibody responses and limited cytokine induction following the first mRNA dose, followed by enhanced humoral and innate cytokine responses after the second dose. No significant differences were observed between the two mRNA platforms despite significant differences in their doses.

### Longitudinal profiling of PBMCs with a single-cell multiome assay

To investigate the cellular dynamics underlying vaccine-induced immune responses, we performed longitudinal profiling of PBMCs from 15 participants who received mRNA-1273 (*n* = 6), BNT162b2 (*n* = 6), or Ad26.COV2.S (*n* = 3) (Fig. 1A) using DNA-Oriented Genome and Transcriptome Mapping by Analysis sequencing (DOGMA-seq). DOGMA-seq is a multi-modal single-cell assay that enables the simultaneous measurement of surface protein expression, gene expression, and chromatin accessibility from the same cells^19^. DOGMA-seq libraries were generated at baseline, day 1, and day 7 for Ad26.COV2.S; at baseline and day 1 after the first mRNA vaccine dose, and at baseline, day 1, and day 7 after the second mRNA dose (Fig. 1A). A total of 69 samples were multiplexed using Cell Hashing^20^ and sequenced in batches of 6-8.

After quality control and filtering, we obtained 429,934 high-quality cells for scRNA-seq and 215,149 for scATAC-seq (Fig. 1D, 1E, and Table S3). Surface protein expression and canonical marker genes identified major immune cell populations including monocytes, natural killer (NK) cells (CD335), B cells (CD19), CD4 T cells (CD4), CD8 T cells (CD8), γδ T cells (TCR-Vδ2), and mucosal-associated invariant T (MAIT) cells (TCR-Vα7.2) (Fig. 1D, and Fig. S1D). Cell type annotations from the RNA-seq modality were transferred to the ATAC-seq modality (Fig. 1E). This dataset enabled a high-resolution, longitudinal analysis of vaccine-induced immune responses, with transcriptional and epigenetic programs captured from the same cells providing direct insights into coordinated regulatory states at single-cell resolution for each vaccine.

### Strong transcriptional interferon responses in myeloid cells after the second mRNA and Ad26.COV2.S vaccines

Clustering of single-cell transcriptomic profiles from the myeloid compartment (*n* = 71,184 cells) revealed three major subsets: CD14 monocytes, CD16 monocytes, and dendritic cells (Fig. 2A, Fig. S2A). One day after Ad26.COV2.S vaccination, the frequency of CD14 monocytes increased substantially, comprising up to 50% of total PBMCs on average, a 2.7-fold expansion relative to baseline (Fig. S2B). No significant changes in cell composition were observed for CD16+ monocytes and DCs. In contrast, mRNA vaccination elicited a more modest but statistically significant increase in CD14 monocytes after both the first dose (p = 0.002, fold-change = 1.25) and the second dose (p = 0.0034, fold-change = 1.72) (Fig. S2B).

**Figure 2.**
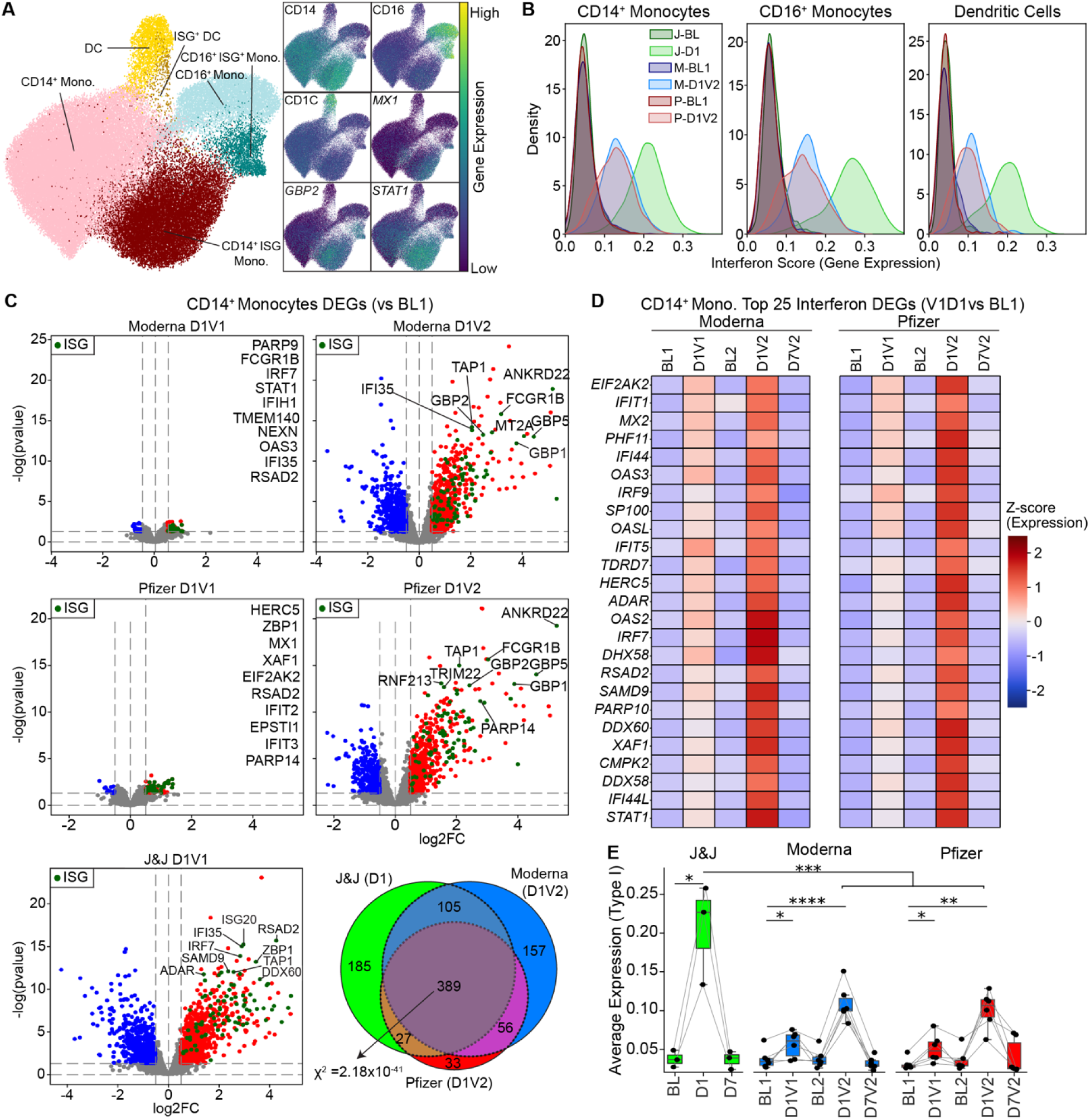
mRNA vaccines induce a robust interferon response after the booster dose. **(A)** UMAP representation of the myeloid compartment from DOGMA-seq, annotated using canonical markers to identify CD14 monocytes, CD16 monocytes, and dendritic cells (DCs). Interferon-stimulated gene (ISG) subsets were defined based on the expression of ISG markers in each subset. **(B)** Kernel density estimation plots of ISG scores (derived from scRNA-seq) for each vaccine and time point show that J&J elicits a strong interferon response one day after vaccination, while mRNA vaccines induce a robust response after the booster dose. **(C)** Differential gene expression analysis in CD14^+^ monocytes comparing post-vaccination time points (D1V1, D1V2) to baseline. Thresholds for significance: log FC > 0.5 and *P* < 0.05. Interferon-stimulated genes (ISG) are highlighted in green. Venn diagram shows the overlap of differentially expressed genes (DEGs) from D1 (J&J), D1V2 (Moderna), and D1V2 (Pfizer). A chi-square test was conducted across the three groups to evaluate whether the overlap of DEGs was statistically significant, indicating a shared transcriptional response. **(D)** Heatmap showing the top 25 interferon-related genes upregulated one day after the booster shot (D1V2) for mRNA vaccine recipients. **(E)** Type-I interferon expression score calculated from manually curated list (n=43), shows stronger Type-I interferon response following J&J vaccination compared to mRNA responses. Statistical significance between timepoints was assessed using two-tailed paired t-test, while comparisons between vaccine groups were performed using the Mann– Whitney U test: n.s. non-significant, \**P* < 0.05, \*\**P* < 0.01, \*\*\**P* < 0.001, \*\*\*\**P* < 0.0001.

Within each myeloid subset, we detected ISG cells characterized by the high expression of interferon-stimulated genes (ISGs), including critical anti-viral and signaling molecules *MX1*, *GBP2*, *WARS*, and *STAT1 (*Fig. 2A and Fig. S2A). These ISG^+^ populations significantly expanded at D1 following Ad26.COV2.S and after the mRNA second dose (Fig. S2C) as confirmed by cumulative ISG scores calculated at the single-cell level using previously published gene set modules (n=96 genes)^21^ (Fig. 2B). Notably, this interferon response was stronger following Ad26.COV2.S compared to the mRNA second dose (p = 0.004).

To further quantify transcriptional changes upon vaccination responses, we performed differential expression (DE) analysis by comparing post-vaccination time points to baseline in CD14 and CD16 monocytes and DCs (Fig. 2C, Table S4). The highest number of differentially expressed genes (DEGs) was detected at D1 after Ad26.COV2.S and at D1 after the second mRNA vaccination in CD14 monocytes (Fig. 2C and Fig. S2C). Across all platforms, 389 genes were commonly induced (Chi-square test, *p* = 3.66 × 10 ^19^) (Fig. 2C), and these were enriched in interferon-associated pathways (Fig. S2D, Table S4), suggesting a shared core interferon response in CD14^+^ monocytes upon mRNA booster vaccination and after Ad26.COV2.S.

Although relatively few DEGs were detected following the first mRNA vaccination in CD14 monocytes, several interferon-stimulated genes (ISGs), including *OAS3*, *IFI44*, and *MX2*, were upregulated (Fig. 2D-E; Fig. S2E). Analysis of curated gene sets (Table S5) revealed that Type I interferon response genes (n=43) were upregulated after the first mRNA vaccine, whereas Type II interferon response genes (n=53) were not (Fig. S2F). In contrast, both Type I and Type II interferon response genes were activated after the second mRNA vaccine and after Ad26.COV2.S vaccination. Notably, Ad26.COV2.S induced a significantly higher Type I interferon response than the mRNA booster, whereas the Type II interferon responses were comparable.

Together, these results show that mRNA vaccination elicits a modest Type I interferon response after the first dose in myeloid cells, followed by a robust, broader interferon program after the second dose. The transcriptional interferon response to Ad26.COV2.S resembled that of the mRNA booster but was markedly stronger in magnitude, driven by the stronger Type I interferon responses.

### The first mRNA dose induces a distinct Type I interferon response state in myeloid cells

We next focused on myeloid cell responses to the first mRNA vaccination by performing a detailed analysis of CD14 monocytes. This analysis revealed three distinct interferon-associated transcriptional states: ISG-low, ISG-dim, and ISG-high (Fig. 3A, S3A). ISG-dim cells selectively expressed a subset of canonical Type I interferon response genes, including antiviral effectors *MX1*, *MX2*, interferon-stimulated genes *IFI44*, *IFI44L*, *ISG15*, and the pattern recognition receptor (PRR) *DDX58/RIG-I* (Fig. 3B, C, Table S6). In contrast, ISG-high cells displayed a broader interferon signature, including the upregulation of IFN-γ–induced guanylate-binding proteins (GBPs), the tRNA synthetase *WARS*, and immunoproteasome components such as *PSMB2* (LMP2)^22,23^ (Fig. 3C). Notably, the ISG-dim state expanded after the first mRNA dose; ∼12% of CD14 monocytes were in this state following BNT162b2 or mRNA-1273 vaccination, compared to ∼5% at baseline (∼2.36-fold increase) (Fig. 3D). This ISG-dim transcriptional signature was restricted to myeloid populations.

**Figure 3.**
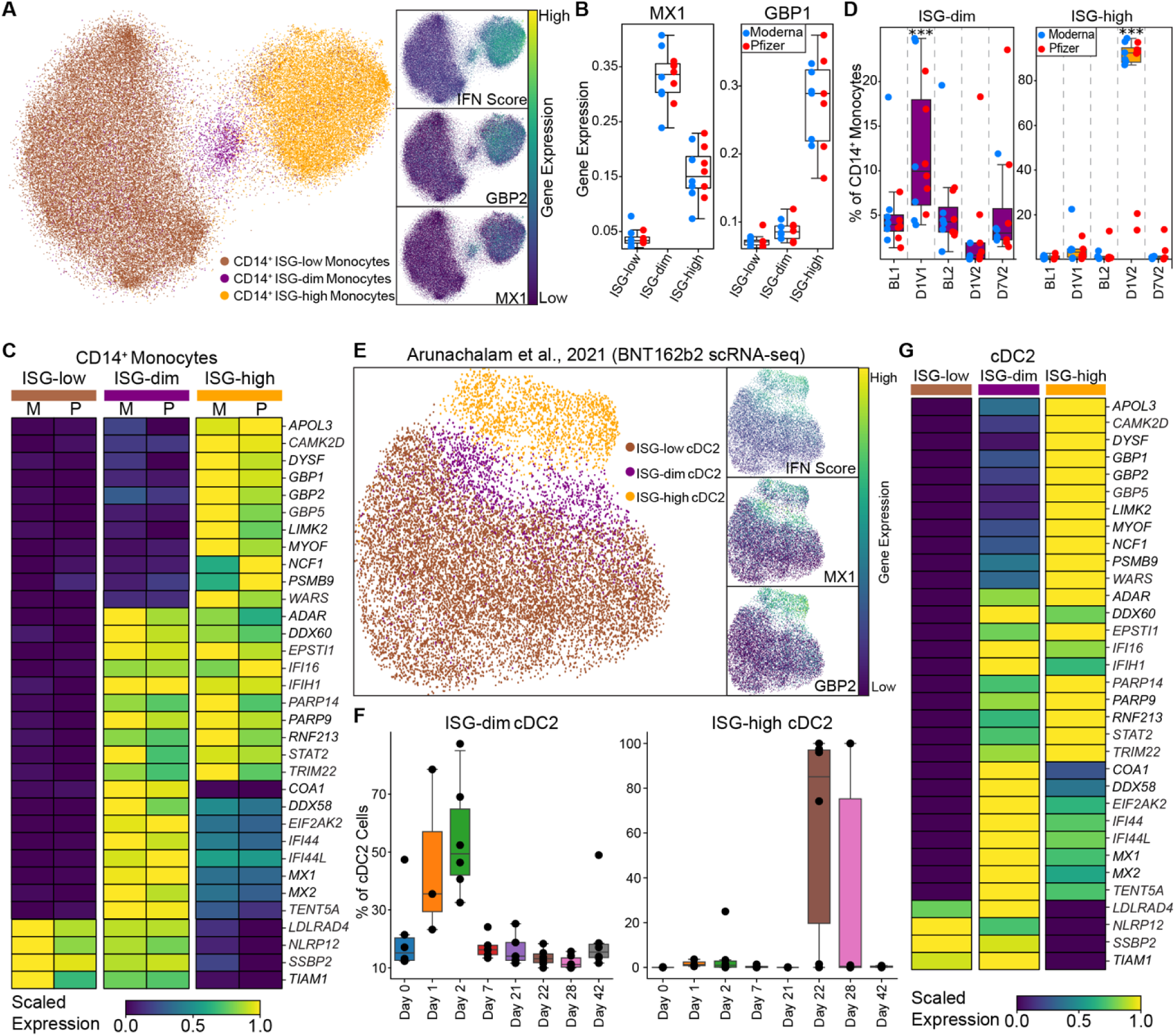
Distinct ISG states associated with primary and booster mRNA vaccination in CD14D monocytes and DCs. **(A)** UMAPs highlights ISG states (ISG-low, ISG-dim, ISG-high) in CD14 monocytes and demonstrate expression of marker gene. **(B)** Box plots show expression of MX1, and GBP1 (**C)** Heatmap showing the expression of genes differentially expressed between three ISG states in Moderna (M) and Pfizer (P) vaccine recipients. **(D)** The proportion of ISG-dim and ISG-high cells within CD14 monocytes across mRNA vaccines and time points. Note that the ISG-dim population expands specifically after the first mRNA vaccination. **(E)** UMAP highlights ISG states (ISG-low, ISG-dim, ISG-high) in cDC2s upon reanalysis of publicly available data. **(F)** The proportion of ISG-dim and ISG-high cells within cDC2s across time points. **(G)** The heatmap displays the expression levels of marker genes for ISG states in cDC2s. **(D)** Statistical comparisons between time points were performed using the one-sided Wilcoxon test: \**P* < 0.05, \*\**P* < 0.01, \*\*\**P* < 0.001, \*\*\*\**P* < 0.0001.

We re-analyzed CITE-seq data from dendritic cell–enriched PBMCs of six BNT162b2-vaccinated individuals^16^ and confirmed the presence of all three ISG states (ISG-low, ISG-dim, ISG-high) in CD14 monocytes in this independent cohort. Furthermore, we detected the same subsets in CD16 monocytes and cDC2s. In all three cell types, ISG-dim state expanded on day 1 after the first vaccine, peaked at day 2, and declined by day 7 (Fig. 3E, Fig. S3B). In contrast, the ISG-high state emerged on day 1 after the second vaccine (Fig. 3F, Fig. S3B, S3C). Marker gene expression in this cohort mirrored our findings: ISG-dim cells highly expressed *MX1*, *MX2*, and *DDX58*, whereas ISG-high cells highly expressed a broader panel of ISGs, including IFN-γ induced genes (Fig. 3G and Fig. S3D).

Taken together, priming induced an early program (ISG-dim state), characterized by Type I interferon response genes, while boosting elicited a robust response (ISG-high state), marked by broader activation of both Type I and Type II interferon response genes.

### Priming induces epigenetic activation of ISGF3 complex in CD14^+^ monocytes

Transcription factors (TFs) such as IRFs (Interferon Regulatory Factor) and STATs (Signal Transducer and Activator of Transcription) are key regulators of interferon responses^24,25^.

To determine whether the ISG-dim and ISG-high states exhibit distinct TF activity, we first examined the ISGF3 complex -the primary effector of type I IFN signaling^26,27^. ISGF3 components, STAT1, STAT2, and IRF9, were upregulated in both ISG-dim and ISG-high CD14^+^ monocytes (Fig. 4A, B). In contrast, IRF1 and IRF8, critical to Type II interferon driven programs^28,29^, were only induced in the ISG-high state (Fig. 4A, B).

**Figure 4.**
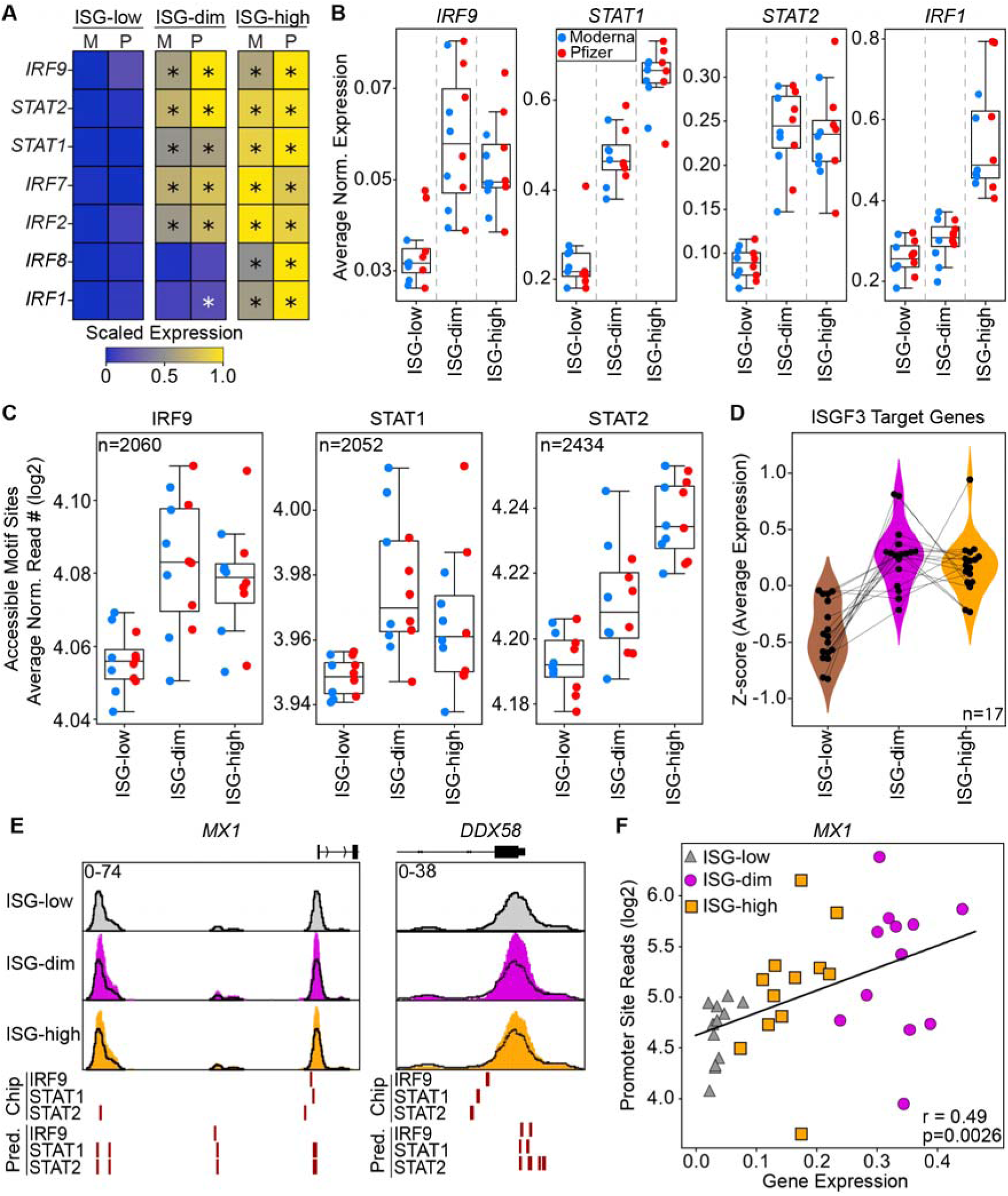
ISGF3 transcription factor complex activity increase in ISG-dim subset. **(A)** Heatmap displaying expression of key interferon response regulatory transcription factors across ISG subsets. **(B)** Average expression of the members of the ISGF3 complex (IRF9, STAT1, and STAT2) and IRF1. **(C)** Average chromatin accessibility at ISGF3 member binding sites across three different ISG subsets using validated binding sites from a published study. **(D)** Z-scores of ISGF3 target genes (n = 17) were calculated based on the average expression of these genes in individual donors across ISG-defined states. **(E)** Genome browsers views of *MX1* and *DDX58* loci across ISG states, highlighting predicted and validated (ChIP-seq) binding sites for *IRF9*, *STAT1*, and *STAT2*. **(F)** Pearson correlation between promoter accessibility and gene expression for *MX1*.

To explore the epigenetic characteristics of these ISG states (Fig. 3A), we analyzed chromatin accessibility in CD14^+^ monocytes. Using previously published ChIP-seq data from human monocytes for STAT1, STAT2, IRF9^26^, we identified validated ISGF3 binding sites^26^ and assessed their accessibility in our ATAC-seq data. ISGF3 binding sites were significantly more accessible in both ISG-dim and ISG-high monocytes compared to ISG-low monocytes (Fig. 4C, Fig. S4A). Only 17 genes had binding sites at their promoters for all three ISGF3 TFs (Fig. 4D, S4B). These genes included *MX1*, *OAS1*, *OAS3*, and *DDX58* (RIG-I), markers of the ISG-dim state. The promoters of these genes exhibited increased chromatin accessibility in ISG-dim cells and harbored ISGF3 binding motifs (Fig. 4 E-F, S4B), further supporting the notion that they are direct ISGF3 targets.

ChromVAR analysis of IRF and STAT motifs, together with longitudinal transcriptomic data, demonstrated that the increased binding accessibility and expression observed for these TFs after the first mRNA vaccine dose is transient (Fig. S5B); no significant changes were detected at the second baseline prior to booster vaccination. Similarly, the changes observed upon booster vaccination (i.e., the ISG-high state) had resolved by day 7 post-vaccination.

These analyses indicate that the transcriptional and epigenetic activation of the ISGF3 complex is a defining feature of the ISG-dim state induced by the priming by mRNA vaccines.

### Exposing monocytes to Type I and Type II interferons recapitulates ISG-dim and ISG-high states, respectively

We hypothesized that Type I and Type II interferons drive the ISG-dim and ISG-high transcriptional states, respectively. Supporting this, the frequency of ISG-high CD14 monocytes correlated strongly with elevated plasma IFN-γ levels following the second mRNA vaccine dose (R=0.952, p=5.38×10, Fig. S5A). To test whether IFN-α and IFN-γ are sufficient to induce the distinct ISG states observed *in vivo*, we stimulated human monocytes for 6 hours with IFN-α, IFN-γ, or both, and performed RNA-seq (Fig. 5A). IFN-α stimulation selectively upregulated ISG-dim markers, including *MX1, DDX58*, and *IFI44*, whereas IFN-γ stimulation upregulated ISG-high markers including *GBP5* and *WARS* (Fig. 5B).

**Figure 5.**
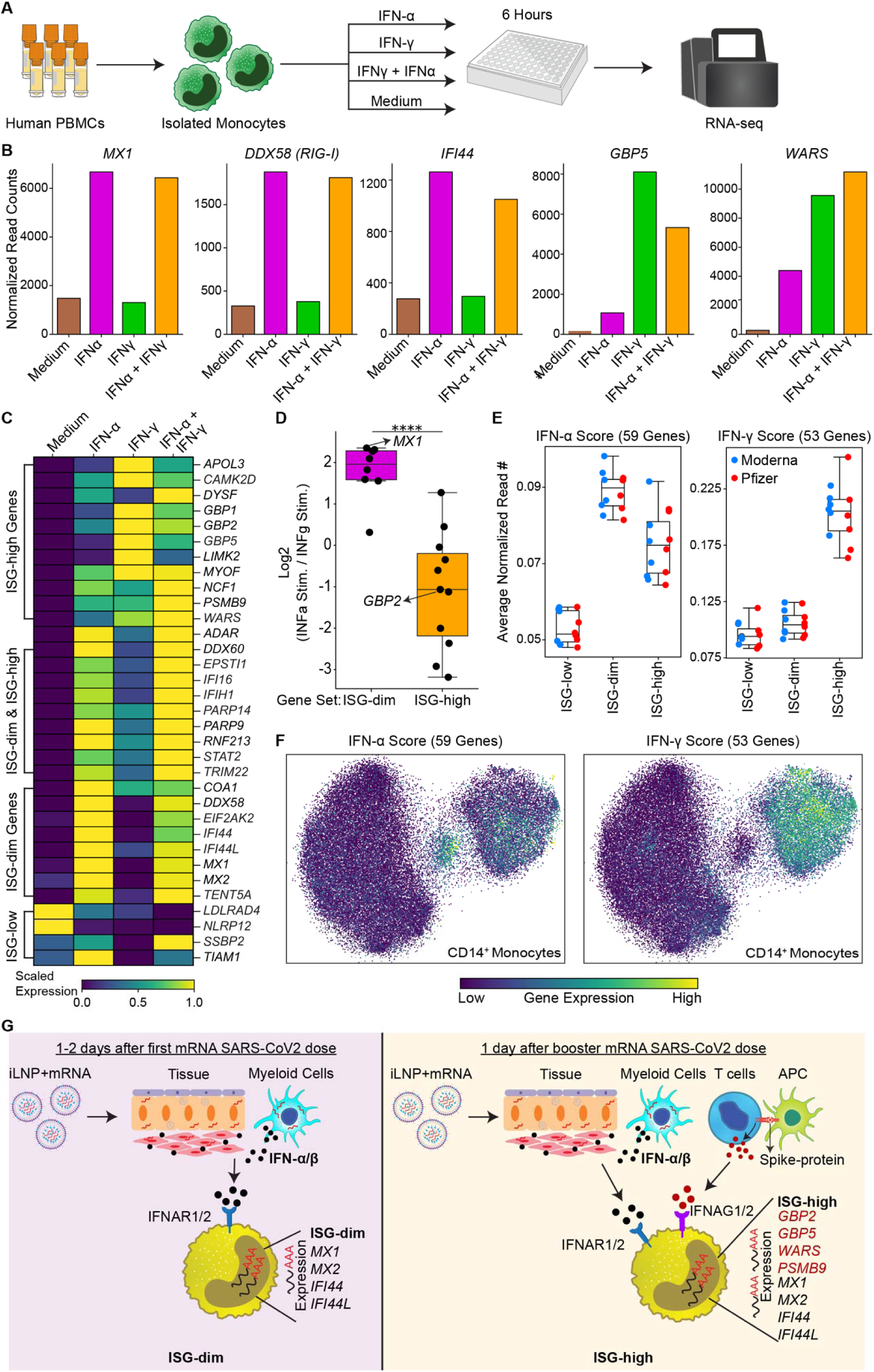
Monocyte stimulation with IFN-α and IFN-γ reproduce *in vivo* ISG states. **(A)** The experimental design involved generating bulk RNA-seq data from isolated monocytes stimulated with either IFN-α or IFN-γ or their combination. **(B)** The expression of *MX1, DDX58, IFI44, GBP5, WARS* upon stimulation. **(C)** The heatmap displays the expression levels of marker genes for ISG states in *in vitro* stimulated data . **(D)** The log fold change between IFN-α and IFN-γ stimulation experiments for markers of ISG-dim or ISG-high states. **(E)** Expression levels of genes in three ISG states for genes associated with IFN-α or IFN-γ stimulation response. Note that ISG-dim state has a high IFN-α score, whereas ISG-high has high IFN-α and IFN-γ scores. **(F)** The UMAP shows the visualization of cells with high IFN-α or high IFN-γ scores. Note the co-localization of high IFN-α cells with the ISG-dim state and co-localization of high IFN-γ score cells with the ISG-high state. **(G)** Summary figure for priming and boosting with mRNA vaccines. The ISG-dim state, driven by Type I interferon expand after the first vaccination and the ISG-high state, that requires both Type I and Type II interferons expand after the second mRNA vaccination. (D) Statistical comparisons were performed using the Mann-Whitney test: \**P* < 0.05, \*\**P* < 0.01, \*\*\**P* < 0.001, \*\*\*\**P* < 0.0001.

Gene expression patterns following IFN-α or IFN-γ treatment closely mirrored the *in vivo* ISG-dim and ISG-high transcriptional profiles, respectively. ISG-dim–specific genes were robustly induced by IFN-α, while ISG-high–specific genes were preferentially activated by IFN- γ (Fig. 5C-D). To further relate these *in vitro* gene programs to *in vivo* single-cell clusters, we derived IFN-α and IFN-γ gene signatures from the stimulation experiments (Fig. 5E, Table S7). UMAP projection of these gene sets revealed distinct localization: ISG-dim monocytes predominantly expressed the IFN-α response gene set, while ISG-high monocytes expressed both IFN-α and IFN-γ response gene sets (Fig. 5F).

Together, these data confirm that Type I interferon signaling is sufficient to drive the early priming response (ISG-dim state) induced by the first mRNA vaccine dose, whereas Type II interferon is required to establish the broader boosting response (ISG-high state) observed after the second dose (Fig. 5G).

## Discussion

In this study, we identified two distinct interferon programs in human myeloid cells (ISG-dim and ISG-high) that are differentially engaged by mRNA priming, mRNA boosting, and adenovirus vaccination. To our knowledge, this is the first head-to-head, single-cell multi-omic comparison of all three FDA-approved COVID-19 vaccines in SARS-CoV-2–naïve individuals. This unique dataset, collected in the unprecedented context of a global pandemic, captures the earliest programming events in human myeloid cells in response to a novel antigen delivered through mRNA and adenovirus vaccine platforms.

Our data reveal that mRNA priming rapidly (1-2 days post-vaccination) induces a transient ISG-dim state in ∼10% of myeloid cells, driven by Type I interferon signaling, whereas boosting and single-dose adenovirus vaccination elicit a more robust ISG-high program involving both Type I and Type II interferons. While prior reports described limited responses after the first mRNA dose^13,15–17^, these have typically focused on bulk transcriptional changes or a single platform. Our work extends these observations by providing cell-type–specific resolution, mechanistic separation of IFN-driven states, and direct comparison across vaccine platforms.

The transcriptional and epigenetic differences between these two states suggests that the innate immune system is differently reprogrammed between priming and boosting. The ISG-dim state was marked by early chromatin accessibility and transcriptional activation of ISGF3 complex components (STAT1, STAT2, IRF9), which in turn regulated canonical Type I interferon response genes and ISG-dim markers *MX1* and *DDX58*. Additional interferon regulatory factors (IRF1, IRF8), typically associated with Type II interferon signaling^30,31^ were activated in the ISG-high state. Although we observed strong early epigenetic remodeling, these changes did not persist: the ISG-dim epigenetic state was not detected at the second pre-boost baseline, in line with prior reports showing limited innate immune memory following mRNA vaccination^17,18,32^. Nevertheless, we cannot rule out the possibility of longer-term reprogramming in bone marrow progenitors^33–35^ or the migration of epigenetically modified myeloid cells to peripheral tissues, either in their existing state or following differentiation into antigen presenting cells such as DCs or macrophages. Moreover, sustained epigenetic changes may require repeated exposures or may be easier to observe in in vitro cultured monocyte-derived macrophages than in freshly isolated blood cells^36^.

*In vitro* stimulation of myeloid cells with IFN-α alone or in combination with IFN-γ reproduced the ISG-dim and ISG-high transcriptional programs induced in vivo, respectively. Type I interferons driving the ISG-dim state might be produced either *via* autocrine signaling by myeloid cells sensing mRNA vaccine components through pattern recognition receptors (e.g., MDA5^9^) or through paracrine signaling from other cell types, such as fibroblasts^37^ or other immune cell types at the site of injection^38–40^. The IFN-γ required to elicit the ISG-high state following booster vaccine is most likely produced by antigen-specific T cells, which expand following booster vaccination^9,12,15,18,41^. Yet, contributions from innate lymphoid cells including, NK cells, cannot be ruled out^42–44^. The precise vaccine component(s) triggering these responses remain unclear. While the mRNA backbone includes nucleoside modifications (e.g., m1Ψ) designed to evade immune recognition, dsRNA contaminants or the lipid nanoparticle formulation may retain immunostimulatory capacity^9,37,45,46^. Future work is needed to dissect which vaccine components drive the early interferon production *in vivo*.

In line with previous studies, mRNA vaccines elicited stronger humoral responses than the single-dose Ad26.COV2.S vaccine, with antibody titers significantly increasing after the second mRNA dose^47–49^. Surprisingly, the Ad26.COV2.S vaccine triggered the most robust interferon response among all platforms, indicating that the magnitude of ISG programs does not necessarily correlate with humoral immunity. It is possible that the memory response to the adenoviral vector^30–32^ diverts the immune response away from SARS-CoV-2 specific targets. The two mRNA vaccines used in our study, BNT162b2 and mRNA-1273, differ substantially in antigen dose (100 µg vs. 30 µg, respectively), yet we observed no major differences in their transcriptional, epigenetic, or antibody responses. Our findings suggest that lower-dose mRNA vaccines may be sufficient to achieve protective immunity.

This study provides the first detailed characterization of myeloid cell responses to priming and boosting with mRNA COVID-19 vaccines, leveraging the unique conditions created by the pandemic, a novel antigen delivered through a novel platform. Our findings underscore the pivotal role of early Type I interferon signaling in shaping downstream immune responses. How these early myeloid programs influence adaptive immunity, particularly in older or immunocompromised individuals, remains an open and clinically relevant question. As mRNA technology continues to expand into vaccines for influenza, RSV, CMV, and other emerging threats, understanding how priming and boosting differentially program innate immunity will be essential for maximizing long-term protection. The framework we establish here can inform the design and clinical evaluation of these new vaccines, ensuring they elicit the most protective interferon-driven responses in diverse populations.

## Limitations

This study offers a single cell-resolution view of mRNA vaccine-induced interferon responses, but several limitations should be acknowledged. First, the cohort size, particularly for the Ad26.COV2.S group was modest, limiting our ability to detect subtle individual or platform-specific differences. Second, while the ISG-dim state emerged as a consistent feature across individuals, its low frequency and transient nature limited our ability to fully resolve its heterogeneity or functional consequences. Third, our focus was restricted to circulating immune cells; missing tissue-resident or lymphoid compartment responses that contribute to early interferon signaling. Finally, while *in vitro* stimulation experiments supported the role of type I and type II interferons in shaping ISG states, they do not fully recapitulate the complexity of *in vivo* cytokine and cellular interactions during vaccination.

## Acknowledgements

We thank the research staff in the UConn Center on Aging and the UConn Older American Independence Pepper Center Program (P30AG067988) for their help in recruitment and sample collection, and the Molecular Profiling team at Immunai for generating the sequencing data. We thank Konstantinos Vlachos, Patrick Petrossian, Mirko Andreoli, and Marisa Mariani in the Molecular Profiling team at Immunai for help with DOGMA-seq analysis. We thank Chew Yee Ngan for the help with bulk RNA-seq data generation. We thank members of the Ucar lab for critical feedback during the study’s progress. This study was made possible by generous financial support from NIH grants under award number AI142086 (D.U., J.B., G.K.). This study was also partly funded by NIH grant U19AI135972 (A.G.-S.) and R01AI160706 (M.S). Opinions, interpretations, conclusions, and recommendations are solely the responsibility of the authors and do not necessarily represent the official views of the National Institutes of Health (NIH).

## Author contributions

J.B., G.A.K., D.U designed the study and raised the funds. G.A.K. and L.K.P. coordinated the clinical sample collection. G.N.E. and Y.Y.Y. led the data analyses. M.S. and A.G.S. generated and interpreted the antibody titer data. E.P.M. generated DOGMA-seq data. R.M. processed blood samples and generated cytokine data. Y.Y.Y., G.N.E, D.U., and J.B. wrote the paper. All authors revised the manuscript and helped with data interpretation.

## Methods

### Study design and cohort

This study was conducted following approval by the UConn Health Center Institutional Review Board (21-149J-1). This prospective, single-site, multi-cohort observational study was designed to determine how vaccine formulations and subject age affect immunologic responses to the SARS-CoV-2 vaccine and to uncover the molecular signatures elicited by novel COVID-19 vaccines. A total of 31 healthy adults aged 21 years and older (21-74 yrs old; mean age 43.7 ± 13.3; 71% female, 29 % CMV) who had received influenza vaccination for the 2020-2021 season but had not yet received a SARS-CoV-2 vaccine were enrolled. Participants who were scheduled to receive an FDA Emergency Use Authorized (EUA) mRNA-based SARS-CoV-2 vaccine as part of the Federal SARS-CoV-2 Vaccination Program were enrolled. After a pre-telephone screening, participants were consented at Baseline Visit 1 or by e-consent prior to the visit.

Blood samples were collected from the participants at eight study visits over a one-year period for mRNA vaccines; BL1 (Day −4 to Day 0 pre-vaccination), D1V1 (1 day after vaccination), D7V1 (Day 7), BL2 (Day 25-28; prior to vaccination dose 2), D1V2 (day after second dose vaccination), D7V2 (7 days after dose 2), D42V2 (Day 70 +/-3days), D154V2 (Day 180 +/-10 days). For Ad26.COV2.S. vaccination, blood samples were collected at six study visit over a one-year; BL (Day −4 to Day 0 pre-vaccination), D1 (1 day after vaccination), D7 (Day 7), D35 (Day 32-35 post-vaccination), D70 (Day 70 +/-3days), D180 (Day 180 +/-10 days). Additionally, blood samples were collected at D365 (∼365 days post-vaccination) for mRNA and Ad26.COV2.S vaccinated participants; however, these samples were not used in the subsequent experiment and analysis.

For all participants, medical history, demographics, vitals (blood pressure, heart rate, temperature), height, weight, BMI, and medication use were collected in the REDCap platform. For individuals 65 years and older, frailty assessments were also performed. For each participant we reported age, sex, ethnicity, and CMV status (Table S1).

### Peripheral blood mononuclear cell isolation

Plasma was separated from the blood collected in ACD tubes by centrifugation, whereas peripheral blood mononuclear cells (PBMCs) were isolated by Lymphoprep (StemCell Technologies) gradient centrifugation within one hour after collection of samples.

### Enzyme-linked immunosorbent assay (ELISA)

For SARS-CoV-2 spike-specific ELISA assays, Nunc MaxiSorp flat-bottom 96-well plates (Invitrogen) were coated with 2μg/mL of recombinant trimeric full-length spike protein (50 μL per well, produced in HEK293T cells using pCAGGS plasmid vector containing Wuhan-Hu-1 Spike glycoprotein gene obtained through (BEI Resources, NR-52394) in bicarbonate buffer overnight at 4°C. Plates were washed three times with 1x PBS + 0.1% v/v Tween-20 (PBST). Subsequently, plates were blocked with 100 μL per well of blocking solution (5% non-fat dry milk in PBST) for 1 hr at room temperature (RT). After removal of blocking buffer, hamster sera were serially diluted in blocking solution starting at 1:100 dilution and incubated for 1.5 h RT. After washing plates with PBST, 50 μL of HRP-conjugated goat anti-human IgG cross-adsorbed antibody (Southern Biotech, Cat 2040-05) was added at 1:5000 dilution. Plates were incubated for 1 hr at RT and washed three times with PBST. Finally, 100 μL tetramethyl benzidine (TMB; BD optiea) substrate was added and incubated at RT until blue color was developed. The reaction was stopped with 50μl 1M H_2_SO_4_, and absorbance was recorded at 450nm and 650nm as a reference. An average of OD_450_ values for blank wells plus three standard deviations was used to set a cutoff value for each plate.

### Determination of Cytomegalovirus (CMV) seropositivity

CMV serostatus was determined by ELISA using the CMV IgG ELISA kits (Aviva Systems Biology). Plasma samples were thawed at room temperature, diluted, and analyzed according to the manufacturer’s recommendations. Calculation of the results was done based on the appropriate controls included in the kit.

### Pilot Sequencing

A pilot sequencing analysis was conducted to determine which visits to focus on and sequence more of. One donor for each vaccine was randomly picked, and eight visits (BL1, D1V1, D7V1, BL2, D1V2, D7V2, D42V2, D154V2) of mRNA recipients and six visits (BL, D1, D7, D35, D70, D180) of the J&J recipient were sequenced and analyzed. Based on this preliminary analysis, five visits (BL1, D1V1, BL2, D1V2, D7V2) of mRNA vaccine recipients and three (BL, D1, D7) of J&J recipients were chosen for sequencing.

### ELLA Microfluidic Immunoassay

The level of four cytokines (IFN-γ, IL-8, IL-10 and TNF-α) and 2 chemokines (CCL2 and CXCL10) in longitudinal plasma samples was quantified using Simple Plex assays run on ELLA microfluidic immunoassay platform (ProteinSimple, San Jose, CA). Plasma samples, diluted 1:1 in sample diluent, were added on the ELLA microfluidic cartridges (human SPCKE-06 cartridge for IL-8/CXCL8, CCL2/JE/MCP-1, IL-10, CXCL10/IP-10/CRG-2, IFN-γ-3rd gen, TNF-α - 2nd gen) according to the manufacturer’s instructions. The ELLA system reported the mean values of the built-in triplicate analysis for each analyte using the manufacturer-calibrated standard curve within 90 minutes.

### DOGMA-seq Library Preparation and Sequencing

Cryopreserved PBMCs were thawed, washed, and stained with barcoded hashing and phenotyping TotalSeqA antibodies (BioLegend) as previously described for DOGMA-seq^19^. Stained cells were fixed in 0.1% formaldehyde for 5 min at room temperature and quenched with glycine solution to a final concentration of 0.125 M before washing twice in PBS via centrifugation at 400 g, 5 min, 4°C. Cells were subsequently treated with the lysis buffer (10 mM Tris-HCl pH 7.4, 10 mM NaCl, 3 mM MgCl_2_, 0.1% NP40, 1% BSA) for 3 min on ice, followed by adding 1 ml of chilled wash buffer (10 mM Tris-HCl pH 7.4, 10 mM NaCl, 3 mM MgCl_2_, 1% BSA). Cells were harvested and diluted in 1× Diluted Nuclei buffer (10x Genomics) and filtered through a 40 µm Flowmi cell strainer before counting using Trypan Blue and a Countess II FL Automated Cell Counter. From there on, the 10x Multiome protocol was followed with the below modifications:

1. During preamplification PCR, 1 μl of 0.2 μM additive primer (ADT and HTO) was added.

ADT add = 5-CCTTGGCACCCGAGAATT*C*C-3

HTO add = 5-GTGACTGGAGTTCAGACGTGTGC*T*C-3

1. 2. 35% of the pre-amplified sample was used to amplify and index protein tags.

GEX, ATAC, ADT, and HTO Libraries were pooled and sequenced on a Novaseq S4 flow cell.

### FASTQ file processing and read alignment

Reads for scATAC-seq and scRNA-seq libraries were mapped and aligned to the reference human genome (GRCh38) using Cell Ranger Arc software (2.0.1). The individual Cell Ranger outputs were aggregated without normalization using Cell Ranger. ADT and HTO reads were processed using kallisto and bustools pipeline^50,51^. Sample demultiplexing was done using HTODemux function from Seurat^52^.

### DOGMA-seq analysis

scRNA-seq, scATAC-seq, and ADT data were processed using Muon and Scanpy libraries^53,54^. Clustering and cell type annotations were done using scRNA and ADT data. scRNA-seq data was total normalized. ADT data was CLR normalized and then total normalized. Highly variable genes were calculated from the scRNA data, and all ADTs except the controls were considered highly variable. After independent normalization, scRNA and ADT data were concatenated, and the rest of the pipeline was performed using highly variable features. After cell type annotations were finalized, only scRNA data were used to discover the ISG^+^ subset. Batch correction was performed with BBKNN^55^. While doing PBMC level annotations, where ADTs are involved in clustering “pool” variable was used as the batch key, and for fine level annotations “library” was used.

scATAC-seq data were processed, and quality control (QC) metrics were assessed using SnapATAC2 from 10x fragment files^56^. Cell multiplets were identified using AMULET with a false discovery rate (FDR) threshold of <5 % and excluded from downstream analyses^57^. Cells with fewer than 1000 fragments or a transcription start site enrichment (TSSe) score below 10 were removed. The intersection of the cells in scATAC data and scRNA data was taken, and the labels of cell subsets were transferred to scATAC data. Peak calling was performed on BAM files for each cell subset using MACS3 with the parameters --call-summits -q 0.05 --nomodel -- extsize 200 --shift -100^58^. The peak matrix for each cell was constructed using SnapATAC2 with a paired-insertion counting strategy^59^.

### Differential Gene Expression Analysis

Three distinct CD14^+^ monocyte subsets were identified ISG-low, ISG-dim, and ISG-high as follows: Mean expression of each gene per donor/subset was calculated, and distributions were compared using two-tailed paired t-test using SciPy library. P values were adjusted using statsmodels library and Benjamini/Hochberg method. Genes that were expressed in more than 10% of cells within any cell subset of interest and expressed in less than 90% of all cells in the analysis were considered in this analysis. The DESeq2 was used to conduct DEG analysis comparing each visit to the corresponding baseline^60^. Prior to analysis, genes expressed in fewer than 1% of cells across all cell subsets of interest were excluded.

### Transcription factor motif binding site accessibility

Iterative overlap peaks were identified using the same MACS3 output for each cell subset, and a peak matrix was generated using SnapATAC2 by extending each peak center by ±250 base pairs (bp). The TOBIAS^61^ (v0.14.0) pipeline was utilized to predict motif binding sites from HOCOMOCO^62^. To identify the experimental binding sites, publicly available IRF9, STAT1, and STAT2 ChIP-Seq data from the THP-1 cell line were utilized^26^. The FASTQ files were processed and analyzed using the MACS3 pipeline^58^. The “bdgpeakcall” function from the MACS3 function was used to call peaks for each TF with a threshold set to yield approximately 100,000 peaks for downstream analysis. Regions present in fewer than 1% of cells across all CD14 ISG subsets that overlapped with ChIP-seq-derived binding sites were selected. ISGF3 binding sites were identified by intersecting IRF9, STAT1, and STAT2 binding sites within scATAC-seq-accessible regions. ChromVAR was employed to assess motif binding site accessibility deviations for each cell subset using HOCOMOCO motifs^63^.

### Cell proportion Analysis

Cell proportions of a subset within a superset were done by simply calculating the percentage of the subset of interest within the superset (all PBMC, the cell type, etc.). Statistical comparisons were done using the one-tailed Wilcoxon test.

### Re-Analysis of Published data

Publicly available dendritic cell-enriched CITE-seq data^16^ was re-analyzed using scanpy. ISG-low, ISG-dim and ISG-high subsets were identified in monocytes and dendritic cells.

### In vitro monocytes stimulation

Human monocytes were purified from PBMCs isolated from healthy donors using Dynabeads untouched human monocyte kit (Thermo Fisher Scientific). Purity of the CD14^+^ cells by flow cytometry analysis was >98%. Purified monocytes, resuspended in complete RPMI medium (RPMI medium 1640 supplemented with l-glutamine, sodium pyruvate, HEPES buffer, penicillin-streptomycin and 10% v/v FBS), were seeded at a density of 1 x 10^6^/mL/well in 96-DeepWell plates (Thermo Fisher Scientific) and stimulated for 6 hours as follows: 100 pg/mL recombinant human IFN-γ (R&D Systems), in the absence or presence of 400 ng/mL of anti-IFN-γ -IgG neutralizing antibody (Invivogen, clone H7WM120), 100 Units/mL recombinant human IFN-α (PBL Assay Science), in the absence or presence of 400 ng/mL of anti-IFN-α-IgG neutralizing antibody (Invivogen, clone H7WM116), 100 pg/mL IFN-γ + 100 Units/ml IFN, or medium control (unstimulated). At the end of incubation, monocytes were collected by centrifugation, and the cell pellets were resuspended in RLT buffer (Qiagen) for total RNA extraction and processing for bulk RNAseq, as described below.

### Bulk RNA sequencing of monocytes and analysis

RNA-seq libraries were prepared with Watchmaker mRNA Library Prep kit (Watchmaker Genomics) according to manufacturer’s instruction. Briefly, 100ng total RNA was used as input. Two ul of 1:1000 dilution ERCC (Thermo Fisher) was spiked into each sample. First, poly A RNA was isolated from total RNA using oligo-dT magnetic beads. Purified RNA was then fragmented at 85C for 10 min, targeting fragments range 300-350bp. Fragmented RNA is reverse transcribed with an incubation of 25C for 10mins, 42C for 15 min and an inactivation step at 70C for 15min. This was followed by second strand synthesis and A-tailing at 42C for 5 min and 62C for 10 min. A-tailed, double stranded cDNA fragments were ligated with xGen™ UDI-UMI Adapters for Illumina (IDT). Adaptor-ligated DNA was purified using KAPA Pure beads (Roche). This is followed by 12 cycles of PCR amplification and another cleaned up. Purified library were sequenced on Nextseq 2000 (Illumina), generating paired end reads of 150bp. Paired-end reads were pre-processed with adapter trimming and quality filtering using Cutadapt^64^. Reads were aligned to the reference transcriptome (GENCODE v38) and quantified using Salmon (v1.10.3) with positional bias corrections and fragment GC bias^65^. The gene level counts were generated using tximport (v1.34.0)^66^. The counts were normalized with TMM using edgeR (v4.4.2)^67^.

### Interferon Stimulated Genes (ISG) score calculations and Density Estimates

Interferon scores were calculated by taking the mean of genes from the interferon modules of BloodGen3Module^21^. The ISG score density estimates were done using kernel density estimates via the Seaborn library’s kdeplot function. In scATAC-seq analysis, genomic regions were prefiltered to retain those present in at least 1% of cells across all analyzed subsets. Smooth quantile normalization was performed using SNAIL^68^ across each donor, followed by log2 transformation of read counts. The regions were annotated to their nearest gene using HOMER^69^ and overlapped with genes from interferon modules from BloodGen3Module. The mean accessibility of the genomic regions was calculated to derive the epigenetic ISG score. Type I and type II interferon scores were calculated using the manually curated list of interferon genes (Table S5). The IFN-α and IFN-γ gene lists were generated based on the fold change between medium and each other (Table S7).

### Statistical Test

Antibody Titer level and Cytokine level comparisons between visits were done using a two-tailed paired t-test. Statistical comparisons of gene expression and chromatin accessibility from scRNA-seq and scATAC-seq were performed using a two-tailed paired t-test between visits at the donor level.

## Data Availability

Raw read data from DOGMA-seq and RNA-seq experiments will be made available on dbGaP upon acceptance of the manuscript (accession number= phs004038). In the meantime, processed data can be viewed and explored at: https://ucarlab.github.io/Covid19VaccineResponses/.

## Code Availability

The scripts used for data processing and figure generation are available on GitHub: https://github.com/UcarLab/Covid19VaccineResponses.

---

**Figure S1:**
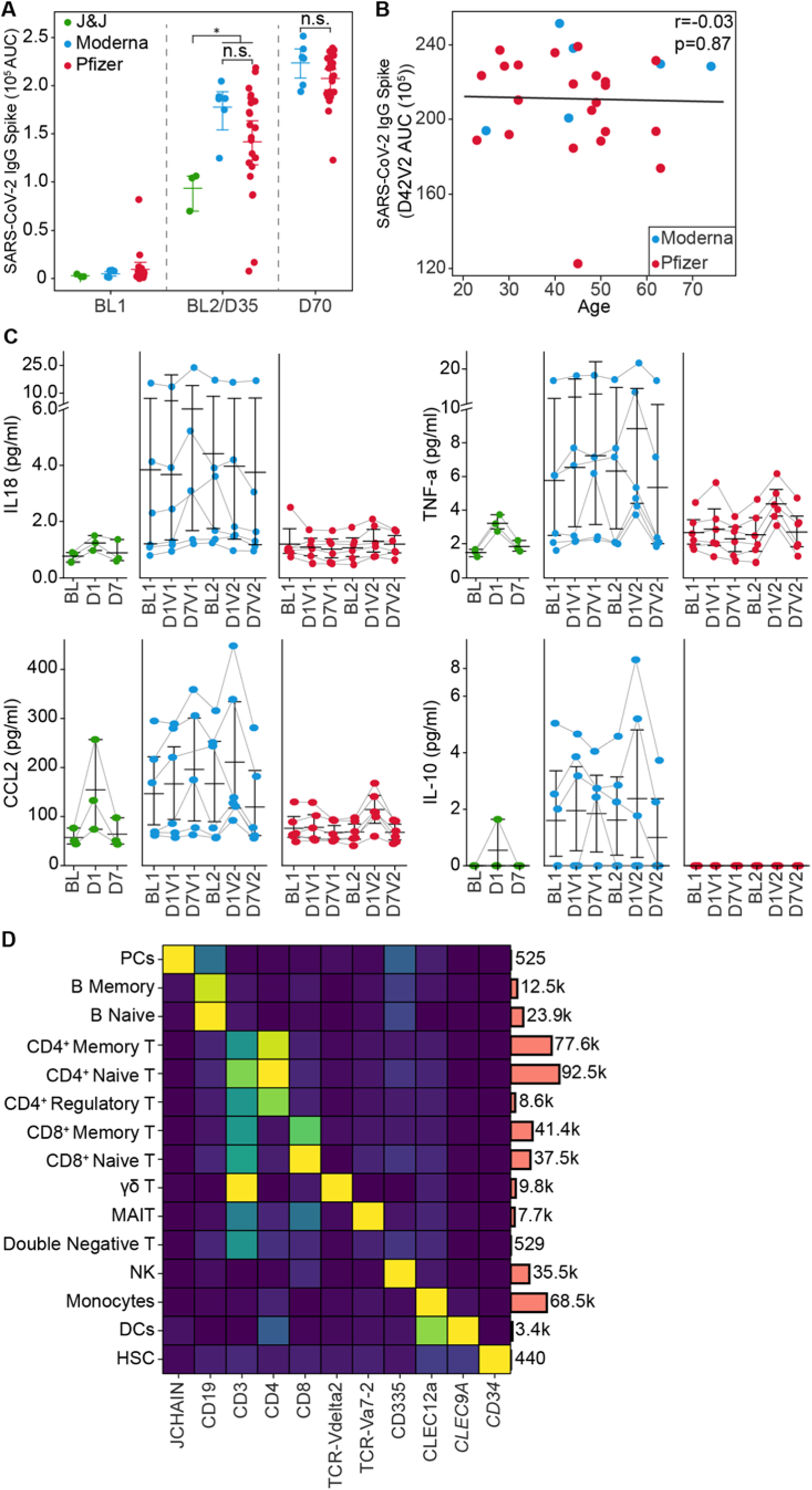
Antibody titer and cytokine level after adenovirus and mRNA vaccination. **(A)** IgG titers against SARS-CoV-2 spike protein measured by ELISA. mRNA vaccines elicit higher antibody titer compared to J&J. (**B)** The correlation plot with age and antibody titer for mRNA vaccines. Pearson’s correlation was used to calculate statistics. **C)** IL-8, TNFα, CCL2, and IL-10 cytokine levels, quantified by ELLA. **D)** The heatmap shows the marker genes for single-cell annotations. (A) Statistical comparisons were performed using the Mann-Whitney test: n.s.: non-significant, \**P* < 0.05, \*\**P* < 0.01, \*\*\**P* < 0.001, \*\*\*\**P* < 0.0001.

**Figure S2:**
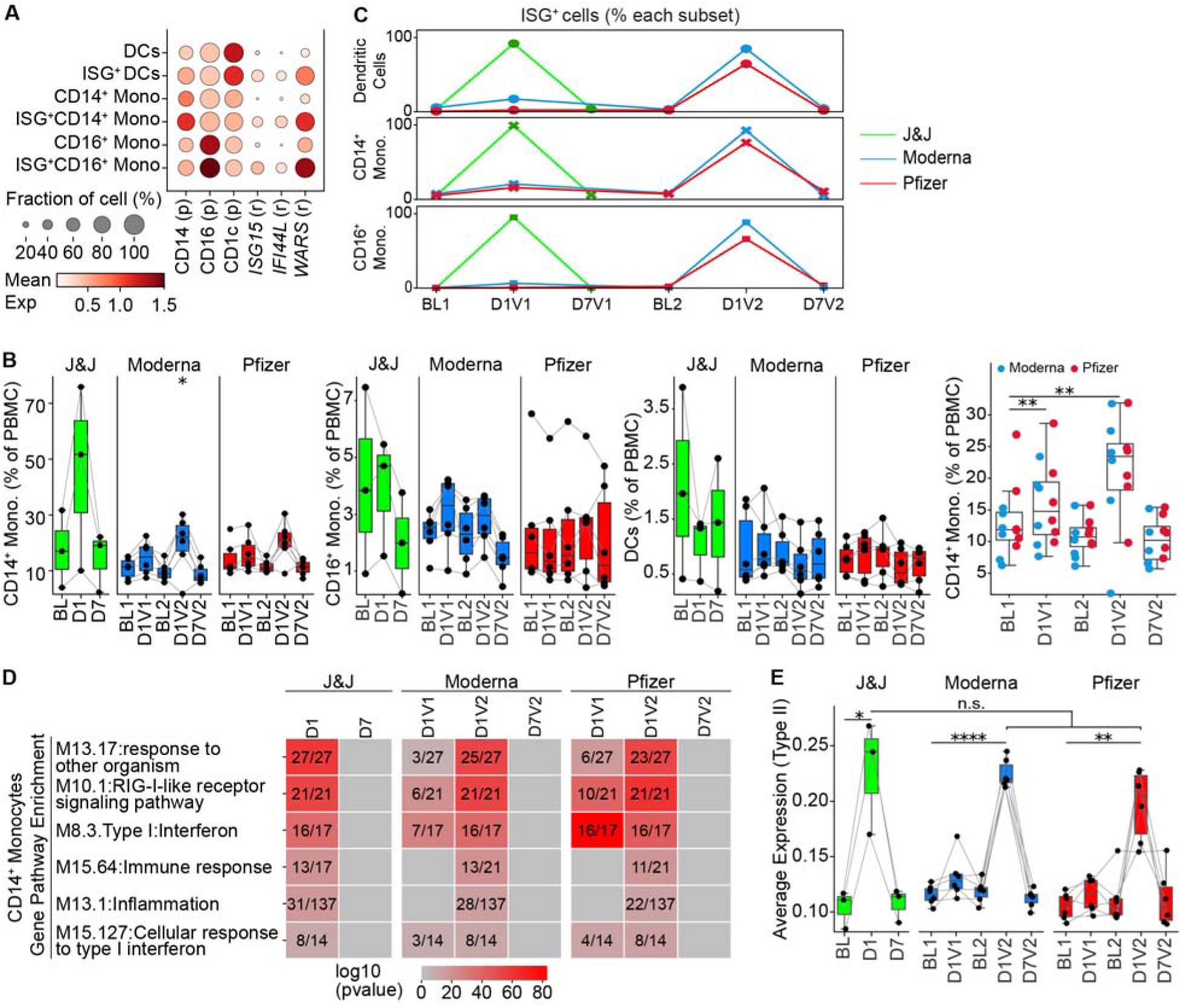
Adenovirus and mRNA vaccines response in CD14^+^ monocytes. **A**) The average expression of marker genes in each myeloid and ISG subsets. **B)** The percentage of CD14^+^ monocytes, CD16^+^ monocytes and dendritic cells (DCs) in total PBMC (left). Significant expansion of the CD14^+^ monocytes upon mRNA vaccination (right). (C) The percentage of ISG subsets within each lineage for each vaccine. **D)** The heatmap shows the top 6 enriched pathways obtained by over-representation analysis using the Blood3GenModule. The number inside indicates the number of overlaps for the respective module. **E)** Type-II interferon expression score calculated from manually curated list (n=51). (B) Statistical comparisons were performed using the one-sided Wilcoxon test, (E) Statistical significance between timepoints were performed using two-tailed paired t-test, statistical significance between vaccine was performed using Mann-Whitney test: n.s. non-significant, \**P* < 0.05, \*\**P* < 0.01, \*\*\**P* < 0.001, \*\*\*\**P* < 0.0001.

**Figure S3:**
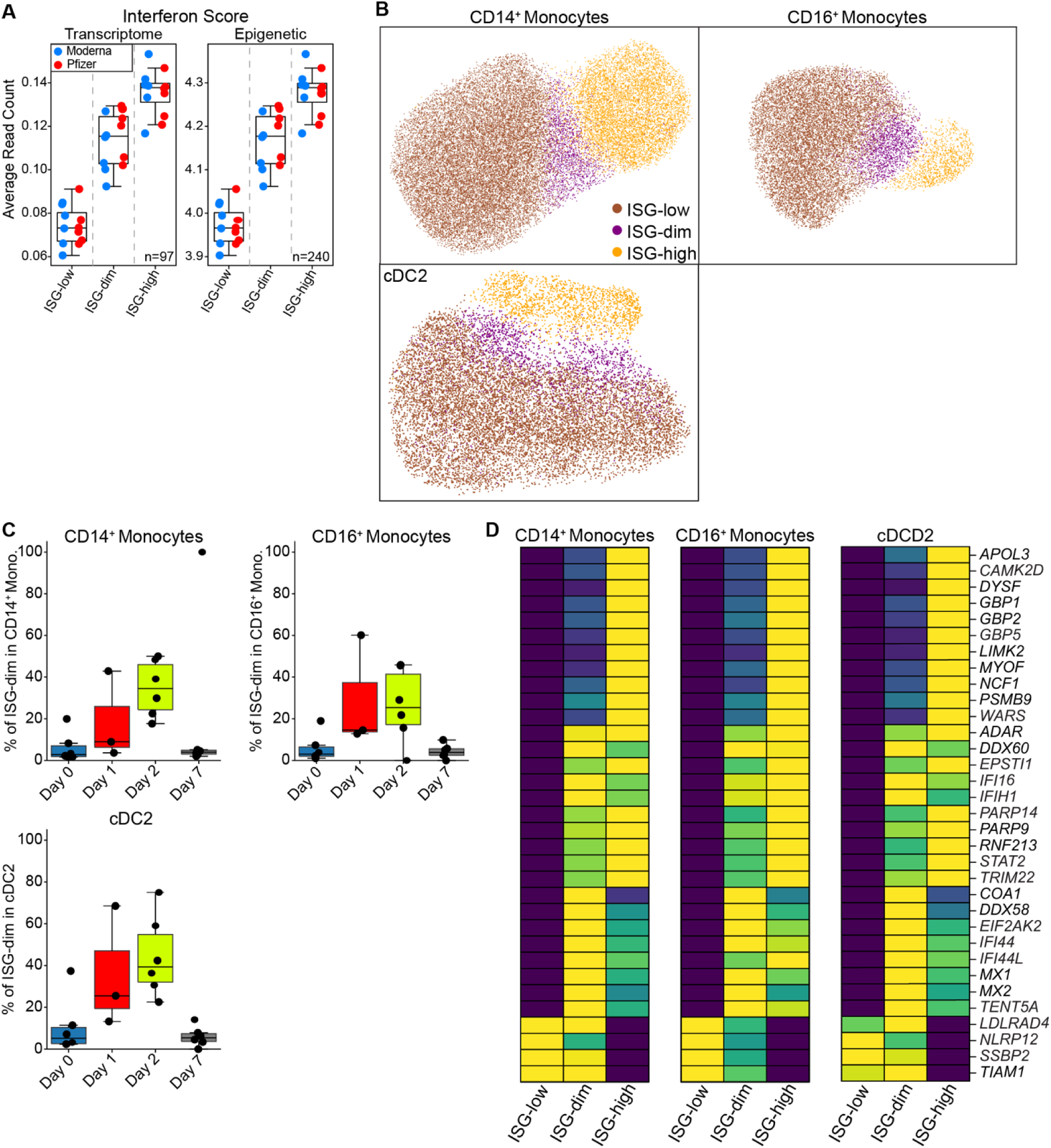
Distinct ISG subset signature in publicly available data from donors vaccinated with BNT162b2. **(A**) Interferon scores for cells in each ISG state. **(B)** UMAP representation of ISG-low, ISG-dim, and ISG-high subsets in CD14^+^ monocytes, CD16^+^ monocytes and cDC2 cell types. (**C)** ISG-dim cell percentage within respective cell populations across different timepoints. **(D)** The heatmap displays the expression levels of marker genes for ISG states in CD14^+^ monocytes, CD16^+^ monocytes, and cDC2s.

**Figure S4:**
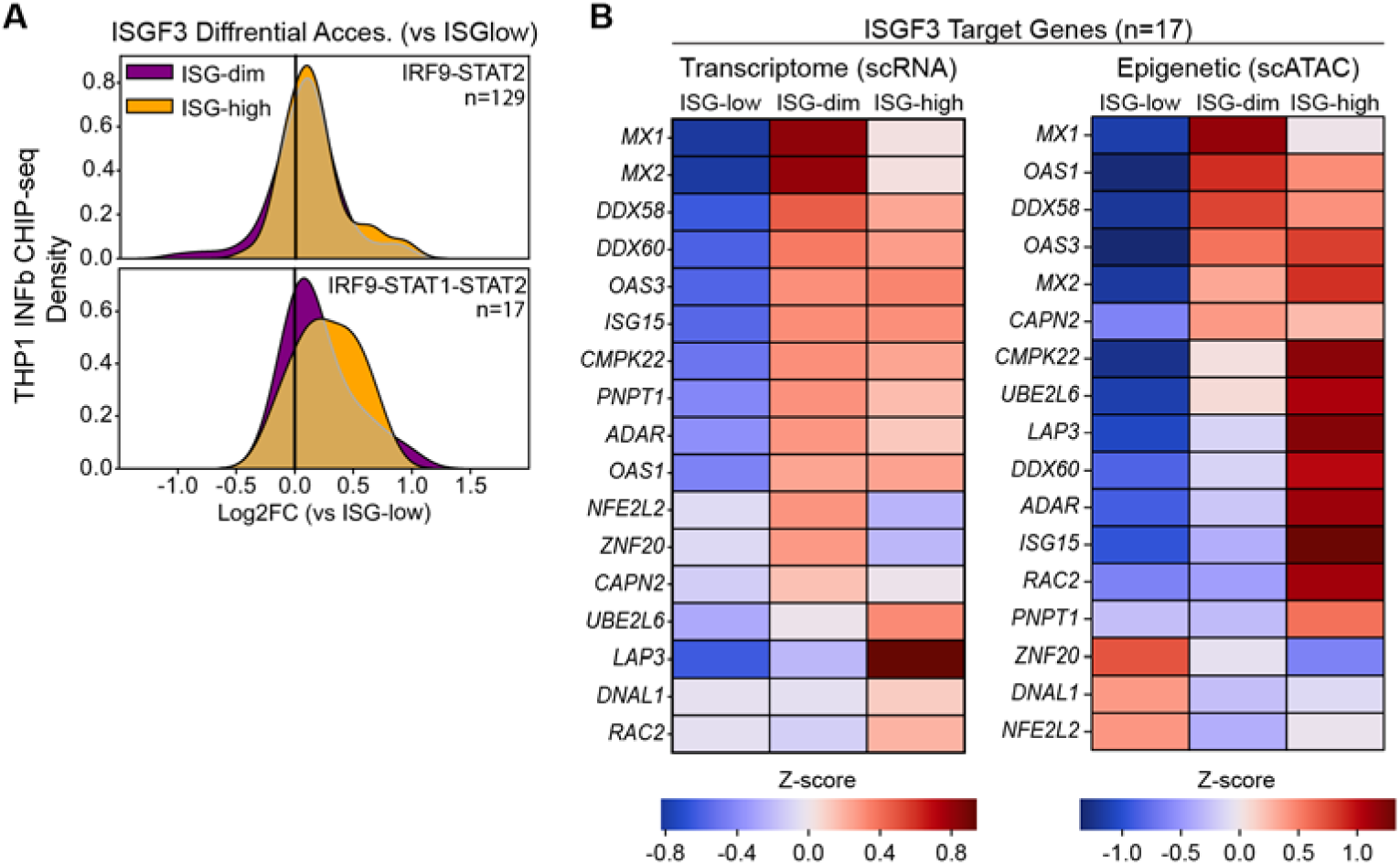
ISGF3 Transcription factor epigenetic dynamic changes. **A**) Kernel density estimation plots show the distribution of accessibility of transcription factor binding sites in ISG-dim and ISG-high compared to the ISG-low subset. IRF9-STAT2 and IRF9-STAT1-STAT2 (ISGF3) peaks were identified by intersecting each TF motif of overlapping peaks of CHIP-seq data. **B)** The heatmap shows the expression and accessibility of the genomic region targeted by the ISGF3 complex in three ISG states.

**Figure S5:**
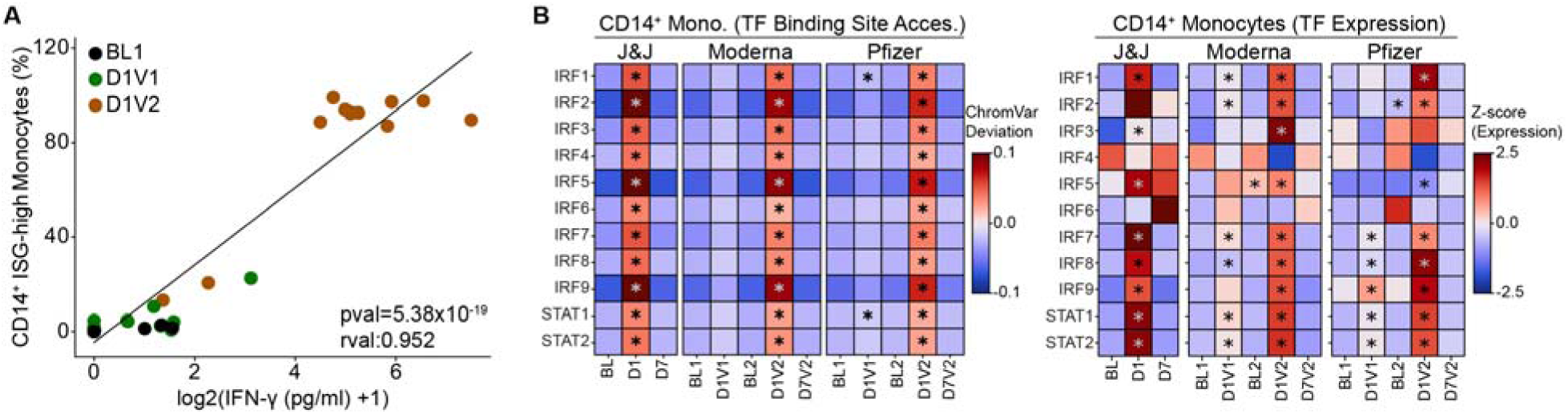
IFNG cytokine levels correlates with ISG-high CD14^+^ Monocyte. **A**) Pearson correlation between ISG-high state percentage in CD14+ monocytes and IFN-γ cytokine levels per sample. **B)** Longitudinal binding site activity of the IRFs and STATs calculated by ChromVAR (left); corresponding gene expression levels (right) of same TFs (right). (B) Statistical significance between timepoints were performed using one-tailed paired t-test, \**P* < 0.05.

## Supplementary Table Legends

**Table S1.** Clinical data for donors vaccinated with mRNA-1273, BNT162b2, and Ad26.COV2.S

**Table S2.** Antibody levels against the SARS-CoV-2 spike protein and concentrations of six plasma cytokines.

**Table S3.** Cell compositional data for all identified cell types across each donor and timepoint from both scRNA-seq and scATAC-seq data.

**Table S4.** The differentially expressed genes for myeloid cells based on the different timepoints

**Table S5.** Manually curated Type I and Type II interferon gene lists.

**Table S6.** The differentially accessible peaks and associated genes in CD14^+^ monocytes for ISG-low, ISG-dim and ISG-high states.

**Table S7.** IFN-α and IFN-γ gene lists inferred from our in-vitro activation experiments.

